# A model of traumatic brain injury using human iPSC-derived cortical brain organoids

**DOI:** 10.1101/2020.07.05.180299

**Authors:** Jesse D. Lai, Joshua E. Berlind, Gabriella Fricklas, Naomi Sta Maria, Russell Jacobs, Violeta Yu, Justin K. Ichida

**Author notes:** These authors contributed equally to the work (Jesse D. Lai, Joshua E. Berlind). **Corresponding Author**, Justin Ichida, PhD, Department of Stem Cell Biology and Regenerative Medicine, University of Southern California, 1425 San Pablo St., BCC 307, Los Angeles, CA 90033.

## Abstract

Traumatic brain injury confers a significant and growing public health burden and represents a major environmental risk factor for dementia. Previous efforts to model traumatic brain injury and elucidate pathologic mechanisms have been hindered by complex interactions between multiple cell types, biophysical, and degenerative properties of the human brain. Here, we use high-intensity focused ultrasound to induce mechanical injury in 3D human pluripotent stem cell-derived cortical organoids to mimic traumatic brain injury *in vitro*. Our results show that mechanically injured organoids recapitulate key hallmarks of traumatic brain injury, phosphorylation of tau and TDP-43, neurodegeneration, and transcriptional programs indicative of energy deficits. We present high-intensity focused ultrasound as a novel, reproducible model of traumatic brain injury in cortical organoids with potential for scalable and temporally-defined mechanistic studies.

## Introduction

Over 60 million new cases of traumatic brain injury (TBI) occur worldwide each year.^1^ Elderly citizens, military personnel, and professional athletes in particular are at elevated risk, with the latter group displaying a 3-fold higher neurodegenerative mortality rate compared to the general population.^2,3^ In severe cases, current pharmacological and surgical interventions fail to significantly improve physical and/or mental impairments, leaving patients with life-long disability.^4^ Previous efforts to identify neurodegenerative mechanisms following TBI have been impaired by the cellular and biophysical complexity of the human brain. Neurons, glia, vascular and immune cells likely all contribute to neurodegeneration through cell-autonomous and non-autonomous mechanisms. Delineating how these proceed individually and synergistically is pivotal to improving our understanding of TBI pathogenesis. Recent studies have linked TBI to pathologic accumulation of the neurotoxic proteins tau, TDP-43 and amyloid-beta, leading to progressive neurodegenerative diseases including chronic traumatic encephalopathy (CTE)^5,6^, amyotrophic lateral sclerosis (ALS)^3,7–9^, Alzheimer’s disease^3,10,11^ and other dementias.^12–14^ Even a single moderate-severe TBI is sufficient to increase risk of developing dementia up to 4-fold.^15,16^ Despite these findings, how primary mechanical injury leads to neurodegeneration and increased risk of dementia-related diseases is unclear. Identifying the interplay between environmental and genetic risk factors for neurodegenerative diseases is critical for the development of therapeutics to mitigate and prevent subsequent pathology.

Current understanding of TBI pathogenesis has been predominantly derived from rodent models, and includes diffuse axonal injury, astrogliosis, metabolic dysfunction, excitotoxicity, inflammation, edema, blood-brain barrier disruption, and hallmarks of neurodegeneration: aggregates of hyperphosphorylated tau and TDP-43.^2,5,6,17,18^ Of these, phosphorylated tau and TDP-43 are particularly common in TBI, and correlate with the extent of neurodegeneration.^5,7,19,20^ In particular, serum phosphorylated tau (Thr231) is reported to be the most accurate biomarker of TBI severity^17^ and is strongly predictive of future neuronal atrophy.^21^ While animal models have proven important for modeling TBI and provide a physical and cellular complexity that relatively mimic humans, many pivotal aspects of neurodegeneration are not faithfully recapitulated, such as substantial differences between human and rodent tau protein.^22,23^ Previous efforts to study TBI in vitro have used stretch and shear forces to either rodent organotypic slices^24^ or rudimentary single- or two-cell type cultures.^25,26^ While these models are useful for studying axonal injury, they have limited translational relevance with regards to the extracellular matrix, stiffness and cell-cell interactions that substantially influence the biophysical properties of mechanical injury *in vivo*.^27–29^ To address these concerns, we adapted a 3-dimensional induced pluripotent stem cell (iPSC)-cortical organoid model^30^ coupled with mechanical injury via high-intensity focused ultrasound (HIFU) with which we can precisely deliver mild/moderate (e.g. sports-related injuries 0.05-0.10 MPa) and severe (e.g. bomb shockwaves >0.6 MPa)^31^ brain injuries.^32^ Organoids generated from this method retain key aspects of human forebrain cellular diversity and cytoarchitecture, are electrophysiologically active and form functional synapses.^30^ Collectively, this system combines a reductionist human cellular model, whereby the cell subsets present can be regulated based on the duration of culture time, together with a highly reproducible and precise mechanical injury platform.

We find that ultrasound injury induces pathologic hallmarks observed in TBI including increased phosphorylated tau (Thr231, p-tau), cytosolic phosphorylated TDP-43 (S409/410, pTDP-43) aggregates and neurodegeneration. Bulk RNA-sequencing (RNAseq) identified changes in hallmark transcriptional programs observed in previous TBI models, including mitochondrial dysfunction and changes in energy production and proteostasis.^2,33,34^ Moreover, we observed dysregulation in the expression and morphology of nuclear pore proteins, which have previously been linked to tau- and TDP-43-related neurodegeneration.^35,36^ These data provide critical support that our ultrasound-induced mechanical injury of iPSC-organoids faithfully recapitulates key features of *in vivo* TBI.

## Results

### High-intensity focused ultrasound induces mechanical injury, neurodegeneration and astrogliosis in human cortical organoids

Human cortical organoids were generated from iPSCs derived from healthy individuals as previously described.^30^ By day 45 of culture, these organoids show substantial neuronal MAP2 expression (Fig 1A) and layered cortical identity through staining with CTIP2 and SATB2 (Fig 1B), and by day 100 of culture develop astrocytes (Fig 1C). Additionally, cortical organoids generated by this method are electrically active, and depolarize in response to glutamate (Supplementary Movie 1).

**Figure 1.**
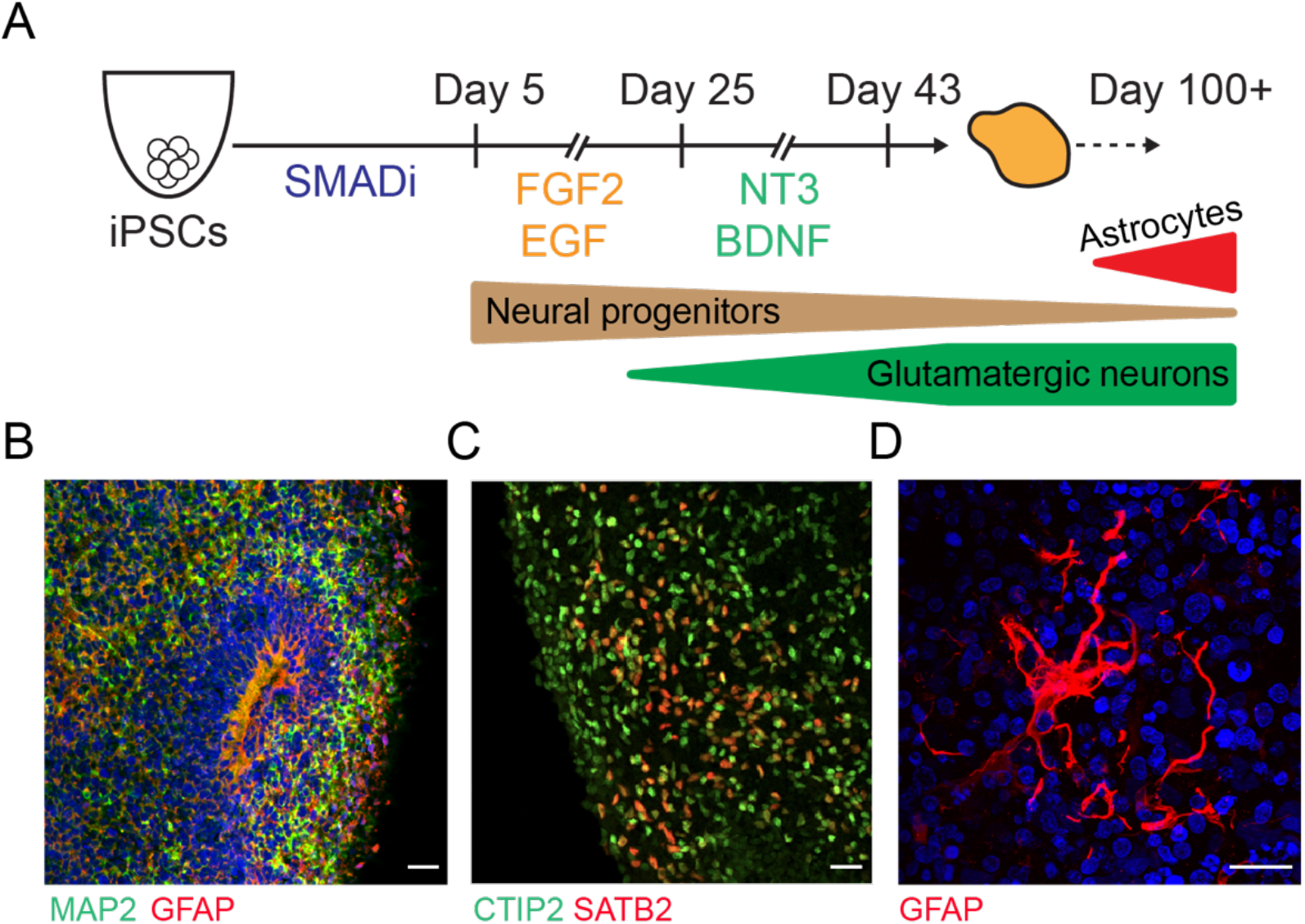
Generation and characterization of iPSC cortical organoids. A) 10,000 iPSCs were resuspended in each well of low-attachment U-bottom plates, and treated for 5 days with 10 μM dorsomorphin and 10 μM SB-431542 for dual SMAD inhibition. Following 20 days of FGF2 (20 ng/mL) and EGF (20 ng/mL), and 18 days of NT3 (20 ng/mL) and BDNF (20 ng/mL) supplementation, organoids will develop neural progenitors, glutamatergic neurons, and astrocytes. B) Immunostaining of day 45 organoids show ventricular zone-like structures containing radial glia (GFAP) and neurons (MAP2). C) Day 45 organoids are positive for deep and superficial cortical markers CTIP2 and SATB2. D) Organoids after 100 days in culture contain highly GFAP positive astrocytes. Images representative of >50 separate differentiations. Scale bars are 50 μm.

We used a custom-built high intensity focused ultrasound that has previously been described and extensively characterized (Fig 2A).^32^ In brief, a waveform generator was used to specifically adjust the average input voltage in millivolts (mVrms) for the transducer. A PVDF hydrophone was then used to detect the acoustic pressure at specified distances from the transducer. A pressure characterization profile was generated based on input voltage and distance (millimeters) from the organoid (Figure 2B). We 3-D printed a conical base scaled to hold an individual organoid in place to ensure the ability to accurately measure the distance from the organoid to the point of acoustic pressure. For the purposes of this study, the ultrasound was set to 510 kHz and applied in 10 ms bursts for two minutes. All injuries, unless otherwise noted, were conducted under these conditions and at an absolute pressure of 0.6 MPa, which mirrors local pressures of a blast injury.^31^

**Figure 2.**
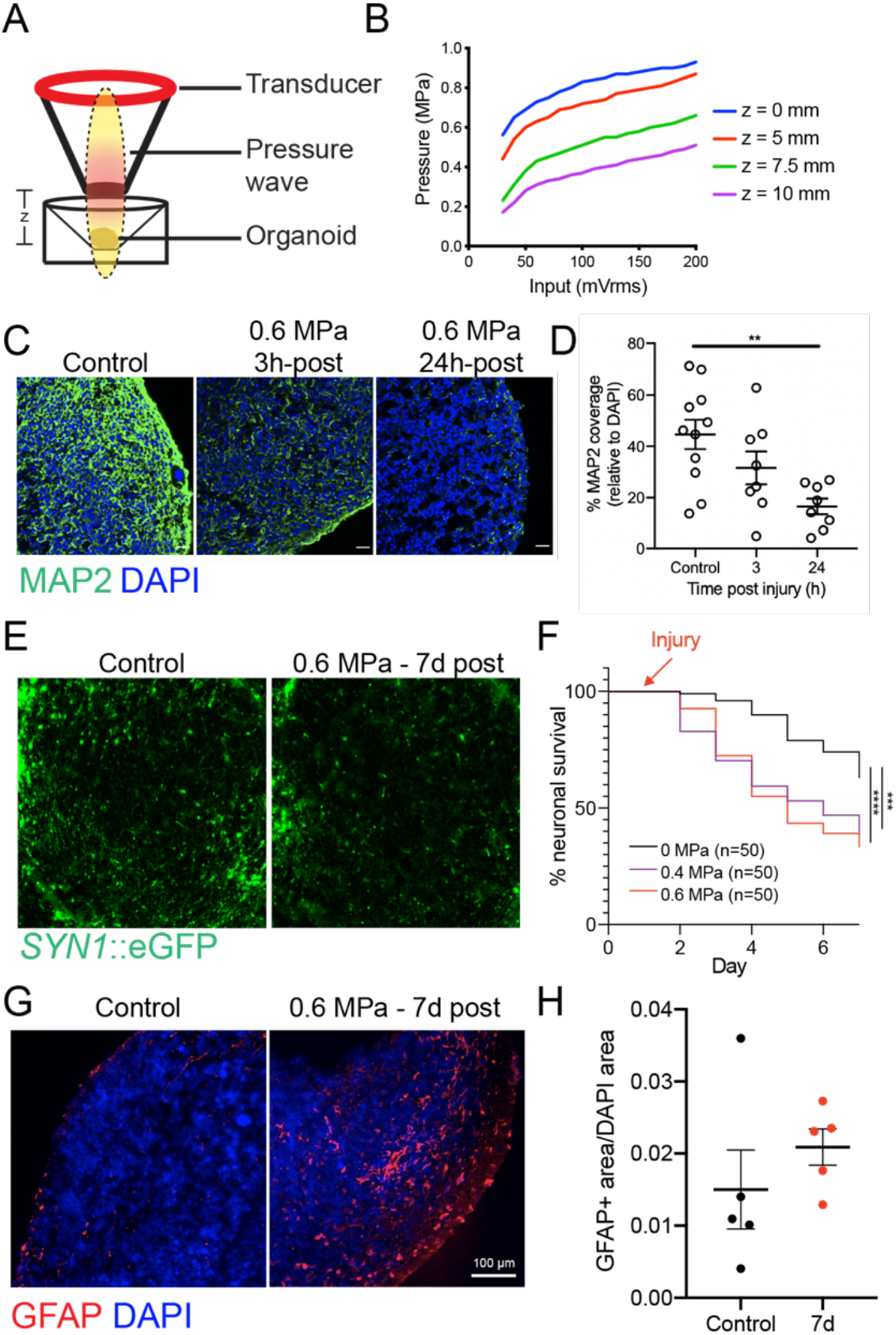
High-intensity focused ultrasound (HIFU) causes neurodegeneration and astrogliosis in cortical brain organoids. A) Simplified schematic of HIFU apparatus which emits an ellipsoid area of acoustic pressure (dotted line). B) The intensity of the pressure can be fine-tuned by adjusting the input voltage and the distance from the organoid. C) A 2 min injury at 0.6 MPa with a 1% duty-cycle shows rapid neurodegeneration in day 45 organoids over 24 hours. D) Cross-sectional quantification of neurodegeneration in immunostained cryosections by area of MAP2 neuronal coverage normalized to the total nuclear area, DAPI. Each point represents 1 organoid, independently injured. E) Day 45 organoids were labelled with a *SYN1*::eGFP lentivirus, and immobilized with Matrigel. An injury of 0.6 MPa as described above was administered directly on the imaging area, and showed marked neurodegeneration over 7 days. F) *SYN1*::eGFP neurons (n=50; 4 organoids) were longitudinally tracked over 7 days. G) Day 100 organoids were injured as described above. Increased GFAP+ astrocytes was observed. H) Quantification of GFAP+ coverage normalized to nuclear area, DAPI. Images representative of at least 5 organoids per condition. Line and error bars represent mean and SEM respectively. Kruskall-Wallis Test and Log-Rank test were used where appropriate. \*\**P*<0.01, \*\*\**P*<0.001, \*\*\*\**P*<0.0001. Unlabelled scale bars are 50 μm.

To determine whether HIFU injury recapitulates neurodegenerative phenotypes observed in TBI, we performed cross-sectional immunofluorescent staining for MAP2 at acute post-injury timepoints. Confocal images show a significant reduction in total neuronal area in injured day 45 organoids compared to mock-injured controls within 24 hours (Fig 2C, D). To confirm this finding, and to eliminate confounding effects of intervening neurogenesis, we measured survival of individual neurons within the organoid by longitudinally tracking lentivirus-labeled *SYN1*::eGFP neurons. This marker of synapsin I-driven GFP expression is selectively expressed in functionally mature neurons, and co-localizes with other neuronal markers including MAP2 (Supplemental Figure 2). Injured and mock-injured control *SYN1*::eGFP organoids were imaged prior to injury and subsequently over 7 days to track survival (Fig 2E). Similar to the reduction of MAP2, we observed a significant, pressure-dependent decrease in neuronal survival in HIFU-injured organoids vs. controls (Fig 2F). These data show that HIFU induces rapid neurodegeneration as expected in a severe TBI model.

Astrogliosis is a common feature observed in post-mortem brain tissues in humans with a history of TBI, and this pattern of astrocyte activation is particularly enhanced around the vasculature.^5^ Although the vasculature is not represented in our organoids, we hypothesized that injury-related molecular signals from neurons and glia would activate astrocytes. Using organoids cultured for 150 days, we inflicted HIFU injury and analyzed GFAP immunoreactivity 7 days after. Similar to post-mortem human samples, we observed increased astrocytic coverage by strong GFAP staining (Fig 2G,H).

### Transcriptional characterization of HIFU-injured organoids

We next performed bulk RNA-sequencing on injured and non-injured organoids to assess global perturbations of major transcriptional networks at 3, 6 and 24 hours post-injury. Principal component analysis showed distinct separation between 24h and control groups (Fig 3A). Differential gene expression from DESeq2 analysis identified 2019 expressed genes by 24 hours of HIFU (Fig 3B). Ingenuity Pathway Analysis (Qiagen) was used to identify the top perturbed canonical pathways (Fig 3C). Consistent with known TBI literature, we found significant changes in several pathways related to energy production^37^ (glycolysis, AMPK Signaling) and protein homeostasis^38–40^ (protein ubiquitination, sumoylation, unfolded protein response) among the top affected pathways.

**Figure 3.**
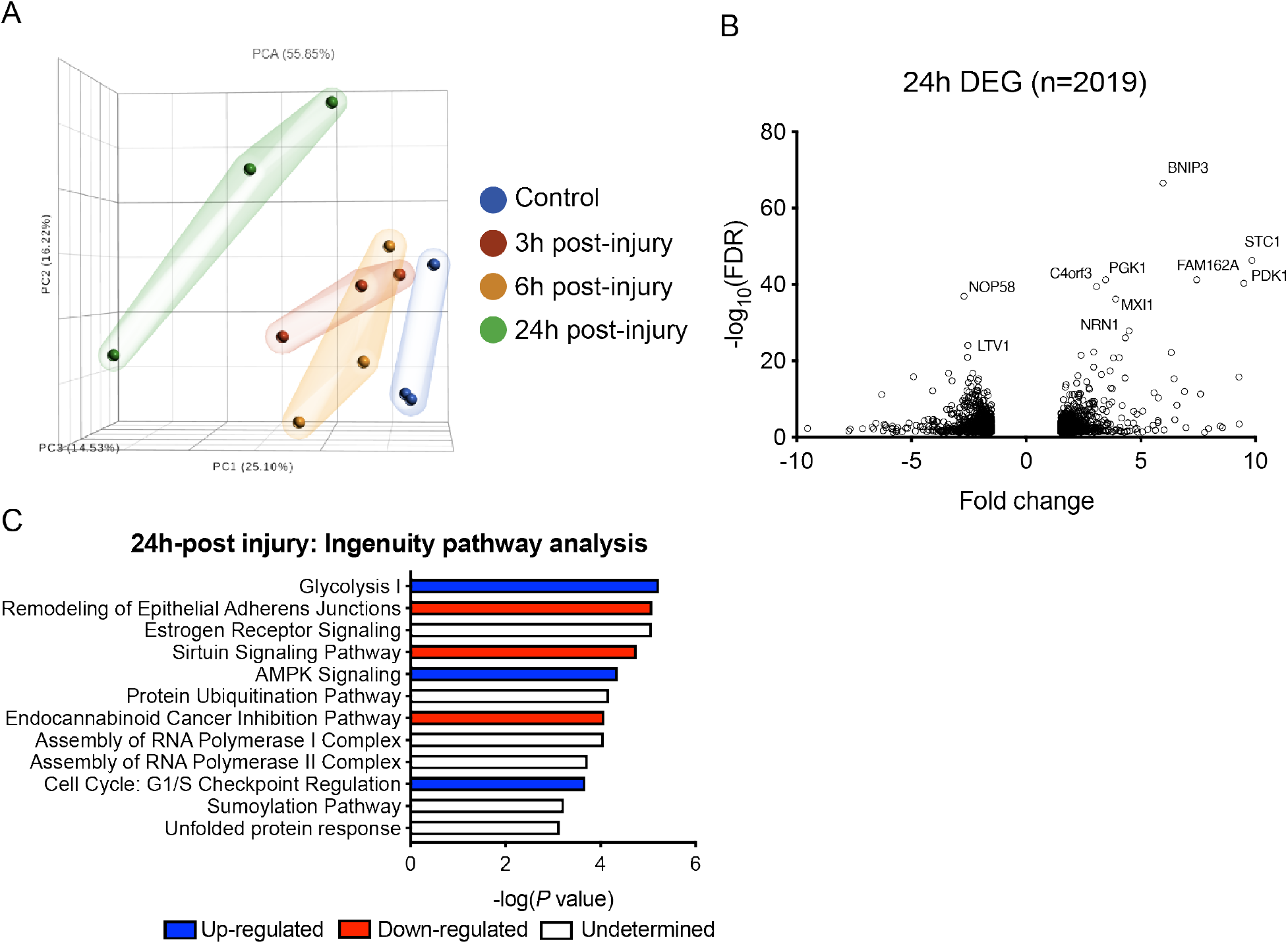
RNA-sequencing of organoids 24h after HIFU-injury. Total mRNA was isolated from day 45 organoids 3h, 6h and 24h after 0.6 MPa (2 min, 1% duty cycle) and prepared for RNA sequencing. A) Principle component analysis comparing each time point. B) Differentially expressed genes (DEGs) in injured organoids compared to control. C) Ingenuity pathway analysis of DEGs 24h post-injury. Three organoids were used in each condition.

### Injured organoids show evidence of phosphorylated TDP-43 and tau pathology

Aberrant aggregation of p-tau is a hallmark of brain injury, and is a key pathological grading criteria for CTE and TBI.^5,41^ Moreover, the ratio of secreted Thr231 p-tau to total tau (p-tau:t-tau) has been shown to be the best biomarker that correlates with TBI severity as indicated by the Glasgow Coma Scale and cranial tomography scans.^17,42^ To this end, we sought to determine whether the p-tau to t-tau ratio (p-tau:t-tau) was elevated in response to HIFU. We collected supernatant from individual organoids at multiple timepoints post-injury and used a multiplexed electrochemiluminescence-linked ELISA to quantitatively assess p-tau:t-tau. Here, we observed a dose-dependent increase in p-tau:t-tau after 0.4 and 0.6 MPa injury (2 min, 1% duty cycle) 3h after injury (Fig 4A). This ratio was significantly increased in organoids derived from multiple healthy control iPSC lines 3 and 24 hours post-HIFU compared to mock-injured controls (Figure 4B,C). We next conducted immunohistological analysis on injured organoids for AT8 (ptau: Ser202/Thr205) and PHF1 tau (ptau: Ser396/Ser404). Interestingly, we observed a decrease in the immunoreactivity 24h post-injury.

**Figure 4.**
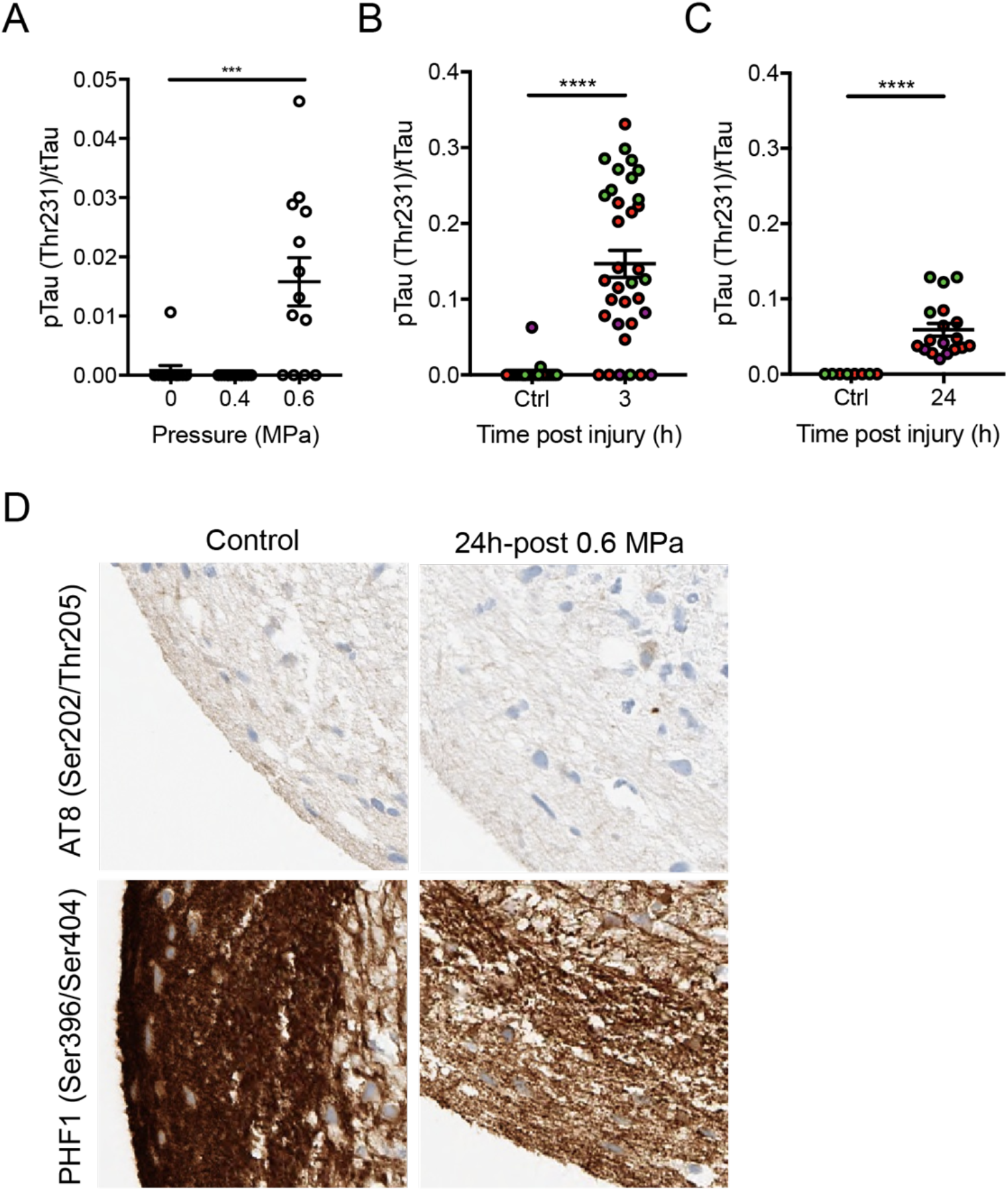
HIFU-injury to cortical brain organoids releases phosphorylated tau. A) Organoids were injured at 0.4 MPa and 0.6 MPa and the ratio of phosphorylated tau (p-tau; Thr231) to total tau (t-tau) in the supernatant were measured by multiplex ELISA 3 hours after injury. The ratio of p-tau:t-tau at B) 3h and C) 24h post injury. Each data point represents 1 independently-injured organoid, each colour represents a different iPSC line. 1 male and 2 female iPSC lines were used in these experiments. D) Representative immunohistochemical staining of formalin-fixed paraffin-embedded organoids with AT8 and PHF1 monoclonal antibodies. Representative of 3-4 organoids per condition. Line and error bars represent mean and SEM, respectively. Kruskal-Wallis test or Mann-Whitney U Test were used were appropriate; \*\*\**P*<0.001, \*\*\*\**P*<0.0001.

Cytoplasmic aggregation of hyperphosphorylated TDP-43 has been shown in ~85% of cases of CTE.^5^ To measure phosphorylation and mis-localization of TDP-43 in our model, we injured day 45 organoids at 0.6 MPa (2 min, 1% duty cycle), and immunostained cryosections for pTDP-43 (S409/410). We observed a coalescence of pTDP-43 puncta and an increase in the intensity of staining 24h after HIFU compared to controls (Fig 5A), however this intensity returned to close baseline over the course of the next 3 days (Fig 5B). We hypothesized that neurons at this younger age have increased plasticity and are more likely to recover from traumatic insults.^43^ In order to test this hypothesis, we injured day 150 organoids with the same parameter described above, and observed sustained cytoplasmic inclusions of pTDP-43 4 days after injury (Fig 5C). After 7 days, these inclusions were not observed, however the intensity of pTDP-43 in both the nucleus (Fig 5D) and cytoplasm (Fig 5E) of neurons remained higher than mock-injured controls.

**Figure 5.**
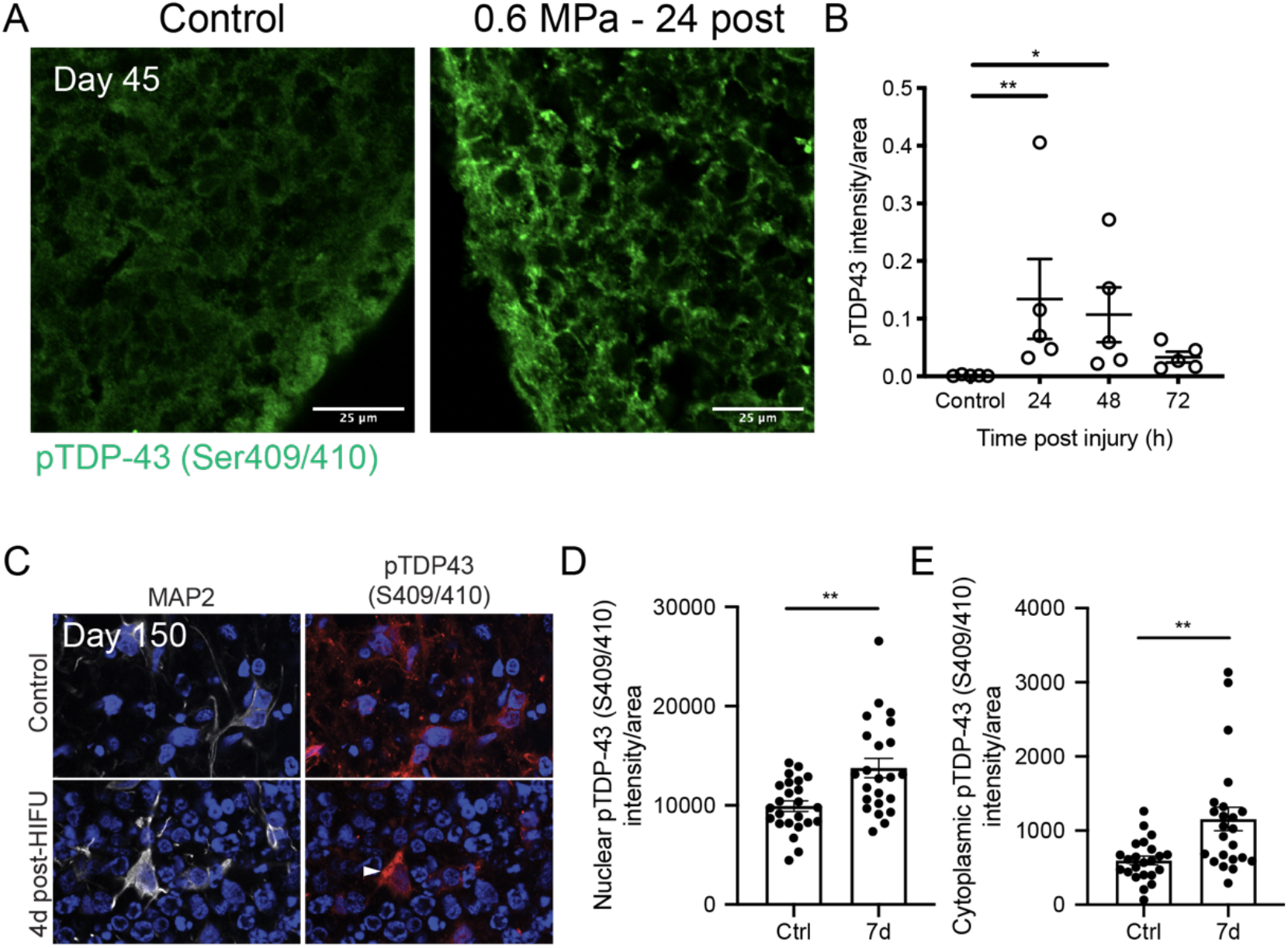
TDP-43 distribution and phosphorylation is altered upon HIFU-injury. A) Day 45 organoids were injured at 0.6 MPa (2 min, 1% duty-cycle) and immunostained for phosphorylated TDP-43 (pTDP-43; Ser409/410) 24h post-injury. B) Quantification of pTDP-43 intensity normalized to total pTDP-43 area over 72h after injury. Each point represents an independently-injured organoid (n=5). C) Day 150 organoids were injured as described above and immunostained for pTDP-43 and MAP2 4-7 days after injury. D) Nuclear and E) cytoplasmic intensity of pTDP-43 was analyzed in MAP2+ neurons 7 days post-injury. Each data point represents 1 neuron across 3-4 organoids. Line and error bars represent mean and SEM, respectively. \**P*<0.05, \*\**P*<0.01 as determined by Kruskall-Wallis Test or Mann-Whitney U Test where appropriate. Scale bars are 25 μm.

### HIFU-injury to organoids disrupt neuronal nuclear pores

From our transcriptomics analysis, we observed significant downregulation of several members of the nucleoporin family after 24h after HIFU-injury, notably the nuclear pore complex proteins Nup98 and Nup62 (Figure 6A). Indeed, nucleocytoplasmic transport deficits have been reported in other neurodegenerative diseases such as ALS, FTD and Alzheimer’s disease, and have been shown to mislocalize and aggregate with tau and TDP-43.^35,36^ Three-dimensional rendering of *SYN1*::eGFP-labeled neurons (Fig 6B) in organoid cryosections revealed a significant reduction in the number (Fig 6C) and volume (Fig 6D) of Nup98-positive pore structures 24h after injury. To directly determine whether HIFU impairs nucleocytoplasmic transport, we used a previously published lentivirus construct, 2xGFP-NES-IRES-2xRFP-NLS, that marks nuclear export with GFP and nuclear import with RFP (Fig 6E).^44^ HIFU injury induced significant loss of nuclear RFP (Fig 4F,G), indicating that nuclear import may be impaired in our model.

**Figure 6.**
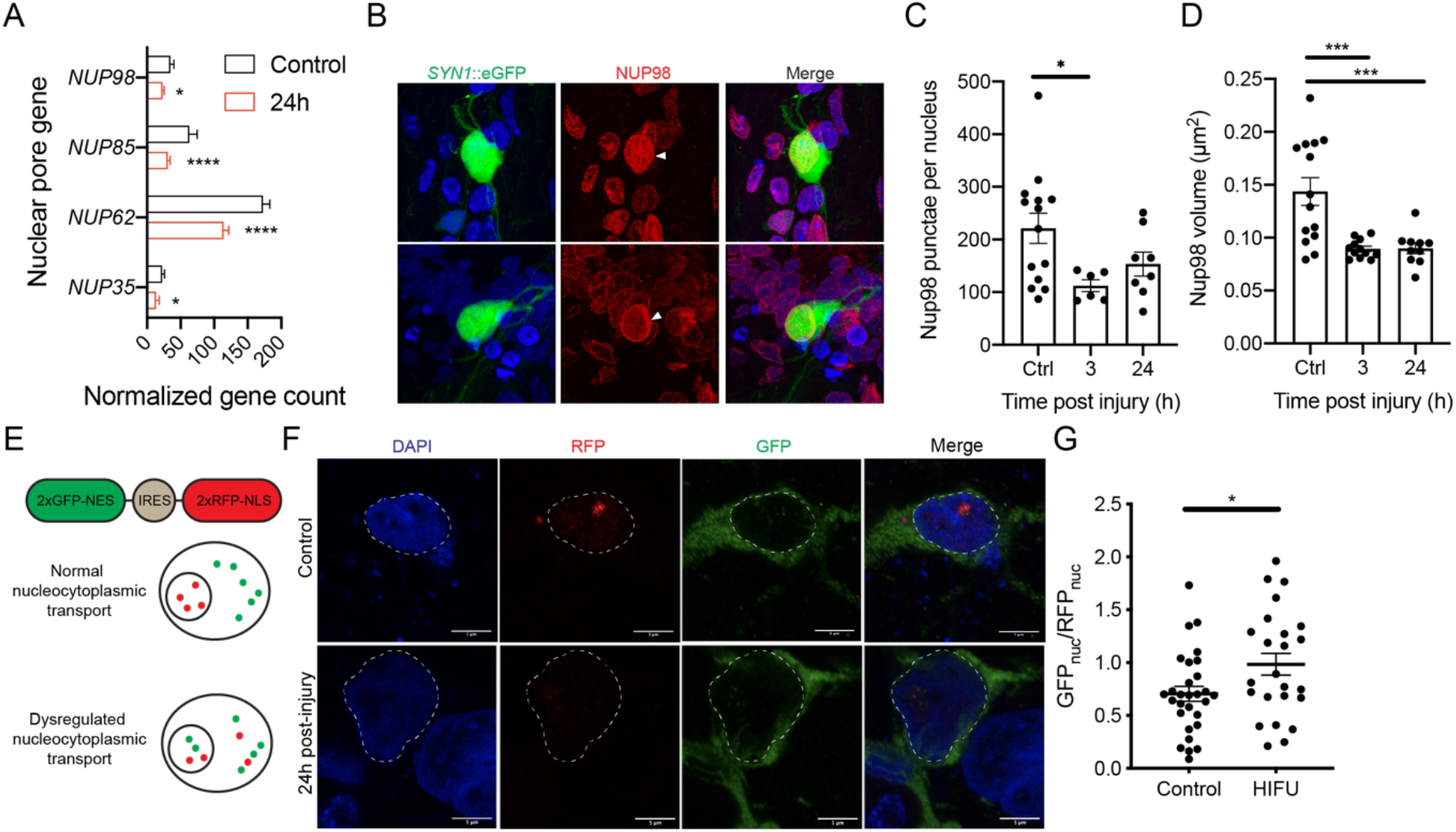
Dysregulation of nuclear pore complexes and nucleocytoplasmic transport after mechanical injury to organoids. Day 45 organoids were injured at 0.6 MPa (2 min, 1% duty cycle). A) Differentially expressed nuclear pore genes 24h after HIFU injury as assessed by bulk RNA-seq. B) Representative immunostaining and quantification of Nup98 C) number and D) volume in *SYN1*::eGFP labelled neurons. Each point represents the average Nup98 volume per neuron across 4-5 organoids per condition. E) Schematic and F) representative images, and G) quantification of nucleocytoplasmic transport deficits and loss of nuclear import of RFP. 25-30 neurons were analyzed across 5 organoids per condition. Line and error bars represent mean and SEM, respectively. \**P*<0.05, \*\**P<0.01*, \*\*\*\**P*<0.0001 as determined by False Discovery Rate, One-Way ANOVA, or Mann Whitney U Test where appropriate. Scale bars are 5 μm.

### Sustained Ca^2+^ influx following HIFU-injury decreases mitochondrial membrane potential and subsequent AMPK signaling contributes to the tau phosphorylation

Another key consequence of TBI is sustained ionic flux (K^+^ efflux, Ca^2+^/Na^+^ influx) that can be caused by events such as ion channel activation, permeabilization of membranes, and acute glutamate release.^45^ In order to compensate for this ion imbalance, mitochondria sequester Ca^2+^ leading to dysfunction and a loss of mitochondrial membrane potential.^46,47^ Here, we observed major changes in genes specific to mitochondrial, including several electron transport chain complex subunits, suggesting that mitochondrial function may be dysregulated (Fig 7A). In order to directly assess whether intracellular Ca2+ increases after HIFU injury, we transduced neurons with a GCaMP6f lentivirus, and observed rapid and sustained increases in intracellular Ca2+ levels immediately after injury in the same neuron (Fig 7B).

**Figure 7.**
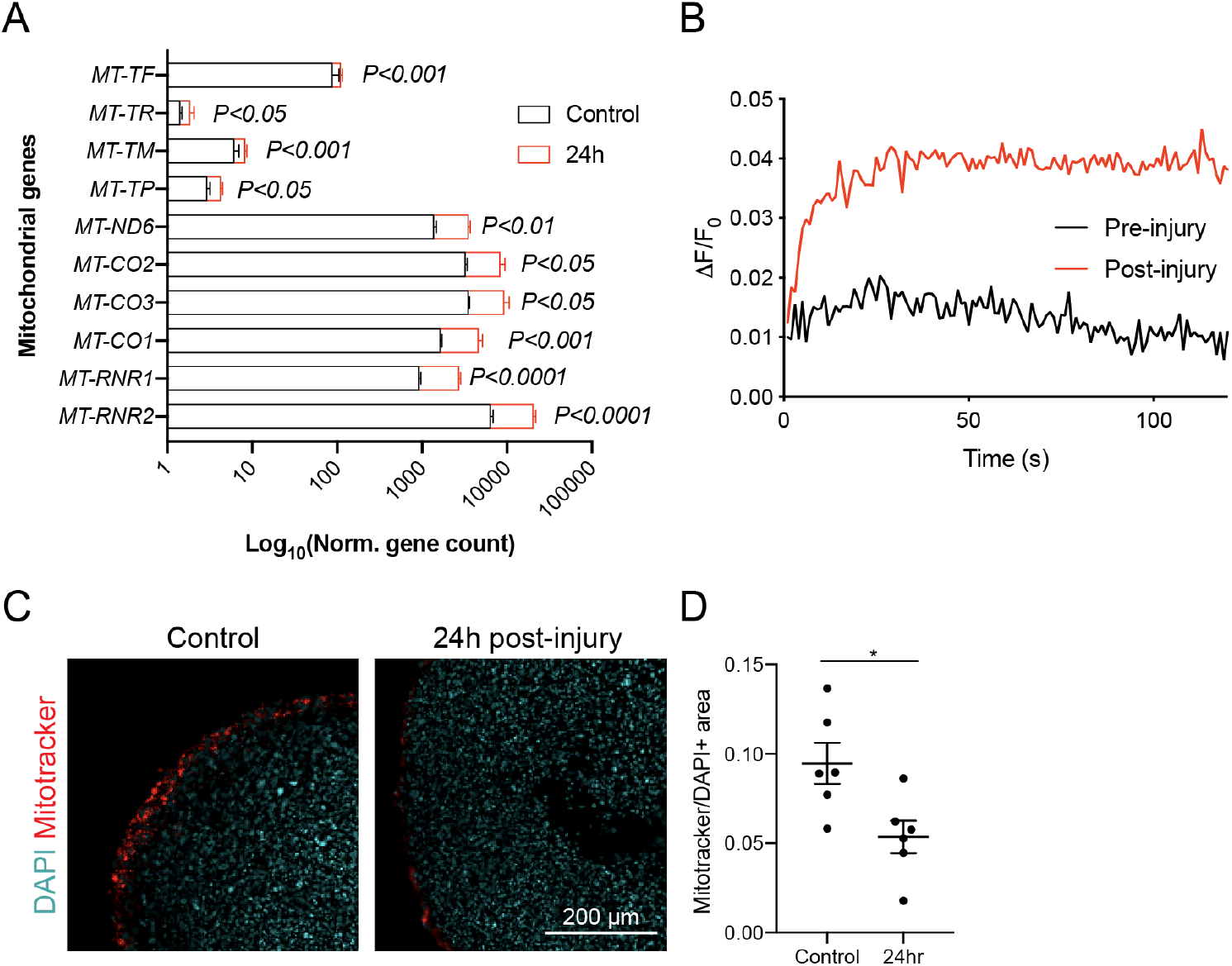
Injury induces sustained Ca^2+^ influx and abolishes mitochondrial membrane potential. A) RNA sequencing show differentially expressed genes in mitochondrial function 24h after injury in day 45 organoids. P values determined by False Discovery Rate. B) Day 45 organoid were labelled with GCaMP6f and fluorescence was recorded pre- and post-injury (0.6 MPa, 2 min, 1% duty cycle) in the same neuron. Representative of 18 neurons from 3 organoids. C) Mitochondrial membrane potential was labelled using Mitotracker CMXRos in day 45 organoids. D) Quantification of CMXRos area compared to nuclear area, DAPI. Line and error bars represent mean and SEM, respectively. Each data point represents 1 independently-injured organoid (n=6). \**P*<0.05 as determined by Mann Whitney U Test.

Saturating the Ca^2+^ capacity of mitochondria leads to dysfunction and a deficit in ATP production.^48^ Our RNAseq data suggest large gene expression changes related to cellular metabolism and energy, specifically increases in glycolysis and AMPK signaling, likely in compensation for low ATP levels (Fig 3B). Indeed, energy crisis is a known disease sequelae following TBI.^34,45^ To assess the global effects of HIFU injury on mitochondria, we measured mitochondrial membrane potential using Mitotracker CMXRos. Here, we observed a significant decrease in the mitochondrial membrane potential 24h after HIFU injury (Figure 7C,D).

## Discussion

We show here that human cerebral organoids generated from iPSCs can be mechanically injured using high-intensity focused ultrasound. In addition to replicating key transcriptional programs that are hallmarks of TBI, including up-regulation of glycolysis and proteastasis, this method accurately recapitulates two key pathological features of neurodegenerative disease: aberrant phosphorylation and accumulation of TDP-43 and tau. Interestingly, in a landmark study of 68 cases of CTE, 37% had co-morbid neurodegenerative disease including motor neuron disease, Parkinson’s disease, Alzheimer’s disease and frontotemporal dementia.^5^ These statistics suggest that TBI plays a significant role in initiation and/or accelerating these pathologies. However, there remains no therapeutic approach that holds promise in reducing this progression to chronic neurodegenerative disease. As a result, the model described here represents and powerful and accurate method to study these mechanisms.

Increased phosphorylation of tau has previously been correlated to neurodegeneration in the context of TBI in a rodent model^41^, and in numerous clinical studies.^5,42^ Similarly, our data suggests a correlation between the intensity of HIFU injury, the amount of ptau secreted in the supernatant and the degree of neurodegeneration. Suppression of pathological tau in mice and non-human primates by monoclonal antibodies or anti-sense oligonucleotides can rescue neurodegeneration, respectively.^41,49,50^ Interestingly, at acute timepoints, our data show an increase in secreted p-tau with a corresponding decrease in p-tau within the organoid. It is possible that this p-tau is derived from dead cells immediately after injury, and may provide an explanation for how tau pathology is transmitted to the contralateral side of the brain following injury as observed in mice.^51^ Indeed, previous studies have shown that, dependent on its phosphorylation, tau is secreted from neurons through direct translocation through the plasma membrane, which can subsequently induce aggregation of the tau repeat domains in adjacent cells.^52^

Aberrant cytoplasmic localization of hyperphosphorylated TDP-43 occurs in >90% of ALS and ~50% of FTD cases.^53^ We show in day 45 cortical organoids, a rapid condensation of pTDP-43 within 24h of HIFU-injury that nearly recovers to baseline over 72h. More mature organoids, cultured for over 100 days, show a more sustained pTDP-43 pathology up to 7 days post-injury, and show hallmark cytoplasmic inclusions as well as pTDP-43 in the nucleus and cytoplasm. Whether or not this is a reflection of the neurons alone, or a contribution from astrocytes is a subject for future studies. Nevertheless, these findings are consistent with previous work that suggest that the developing brain is more resilient to formation of toxic aggregates.^54^ Similar to our data, in 6 patients who died within 3 days following severe TBI, the only TDP-43 pathology observed was an increase of pTDP-43 (S409/410) fragments in the nuclear fraction of the frontal cortex.^55^ Interestingly, this study also showed a TBI-mediated exacerbation of TDP-43 pathology and cell death in a transgenic mouse expressing mutant TDP-43 (A315T), however, significant differences in TDP-43 pathologies between mouse and human samples were observed, thus necessitating the study of TBI in human cells.^55^

Many mechanisms underlying how phosphorylation and dysregulation of tau and TDP-43 lead to neurodegenerative disease have been proposed. In humans and animal models of ALS and FTD, tau and TDP-43 co-aggregate with nuclear pore complex proteins leading to dysregulation of nucleocytoplasmic transport, which can subsequently be rescued by pharmacological inhibition of nuclear export.^35,56^ We show here that the nuclear pore protein, Nup98, is disrupted in neurons within 3h after HIFU-injury. These data suggest that TBI may elicit similar pathways that contribute to the increased risk for neurodegenerative disease.

A key pathological event observed in our model is the influx of calcium leading to mitochondrial dysfunction and energy crisis following TBI.^45^ Previous studies in ALS and FTD suggest that phosphorylation and dysregulation of tau and TDP-43 likely exacerbate these pathological cascades. Indeed, TDP-43 proteinopathy in post-mortem human samples have been shown to localize to mitochondria and impair function, while inhibition of this trafficking rescues neurodegeneration.^57^ Moreover, tau and the its regulation by phosphorylation affects mitochondrial transport^58,59^, dynamics (fission, length)^60^, and function.^61,62^ Mice deficient in tau are protected from excitotoxic neurodegeneration, while TDP-43 alone can elicit similar effects as ion sequestration in mitochondria.^57,63^ Taken together, preventing mitochondria dysfunction may prove to be an effective therapeutic approach for mitigating traumatic brain injury and its contributions to chronic neurodegenerative disease.

Traumatic brain injury is an incredibly complex series of events to which there are no viable treatment options. Moreover, how these events contribute to chronic neurodegenerative disease is largely unknown. The model described here highlights the hallmark disease processes involved in acute TBI, as well as mechanisms that have been shown to influence long-term neurodegeneration. This high-fidelity TBI model bridges the gap between traditional *in vitro* systems and complex higher organisms thereby providing a scalable and genetically-flexible system to identify disease mechanisms and potential therapies for the acute and chronic effects of TBI.

## Author contributions

JDL and JEB designed and performed experiments, analyzed and interpreted data, and wrote the manuscript. GF and NSM performed experiments and analyzed data. NSM and RJ built ultrasound equipment and interpreted data. JKI designed experiments, interpreted data, and edited the manuscript.

## Acknowledgements

This work was supported by Amgen, DoD GRANT12907280, the New York Stem Cell Foundation, the Tau Consortium, NINDS R01 1R01NS097850-01, the Harrington Discovery Institute, the Alzheimer’s Drug Discovery Foundation, the Association for Frontotemporal Dementia, and the John Douglas French Alzheimer’s Foundation. We thank Seth Ruffins and the Optical Imaging Core for imaging support. We thank Yibu Chen and the Norris Medical Library for assistance in analyzing RNA-sequencing data. We also thank Kristen Whitney and John Crary (Icahn School of Medicine at Mount Sinai) for processing of paraffin-embedded organoid histology samples. JKI is a New York Stem Cell Foundation-Robertson Investigator, a John Douglas French Alzheimer’s Foundation Associate Professor, and a USC Merkin Scholar. JDL is an employee of Amgen. JKI is a co-founder of AcuraStem.

Address correspondence to: Justin K. Ichida, 1425 San Pablo Street, Los Angeles, California 90033, USA. Phone: 323-442-0063.

## Methods

### Human iPSCs: culture and CRISPR/Cas9 genome editing

Lymphocytes from healthy donors were obtained from the NINDS Biorepository at the Coriell Institute and reprogrammed into iPSCs as previously described.^29^ Cell lines derived from FTLD-Tau patients harbouring the V337M point mutation, and their corresponding CRISPR-corrected isogenic control, were acquired and generated by the Tau Consortium Stem Cell Group as previously described.^30^ Cells were maintained on Matrigel (BD) in mTeSR1 medium (Stem Cell Technologies).

### Generation of cortical organoids

The generation of human cortical organoids was adapted from a previously published protocol.^19^ In brief, iPSCs at ~70% confluence were dissociated into single cells with Accutase (Stem Cell Technologies) and 10,000 cells/well were seeded into a 96-well U-Bottom Low-Attachment plate (Corning) in mTeSR1 + 10 μM ROCK inhibitor (Y-27632; Tocris) to generate iPSC spheroids. Media was replaced 24h after seeding with mTeSR1 without ROCK inhibitor. For the following 5 days, neural induction was initiated with 10 μM dorsomorphin (Cayman Chemicals) and 10 μM SB-431542 (Cayman Chemicals) in DMEM/F12 containing 20% KnockOut Serum (Gibco), 1 mM non-essential amino acids (Gibco), 1X GlutaMAX and 0.1 mM b-mercaptoethanol. From days 6-15, media was refreshed daily with neural medium (Neurobasal-A (Invitrogen), B-27 Supplement without vitamin A (Gibco), 1X GlutaMAX, 2% penn/strep) supplemented with 20 ng/mL bFGF (Peprotech) and 20 ng/mL EGF (Peprotech), and every other day from days 16-25. On day 20, organoids were transferred to 6-well low attachment plates (Corning) and placed on an orbital shaker rotating at 60 rpm. From days 26-43, media was refreshed every other day with neural medium containing 20 ng/mL NT-3 (Peprotech) and 20 ng/mL BDNF (R&D). Organoids were maintained from day 43 onwards in neural medium and refreshed every 3-4 days.

### Vectors and Viral transduction

Virus was produced by polyethylenimine (PEI; Sigma-Aldrich) transfection of 89-90% confluent HEK293T cells with the viral vector together with the lentiviral packaging plasmids: pPAX2 and VSVG. Media was changed 24h after transfection, and supernatant was collected 48h and 72h hours after transfection. Virus supernatant was passed through 0.45 μm filters, concentrated overnight at 4C with Lenti-X (Clontech) as per manufacturer’s protocol, resuspended in DMEM, and stored at −80C. Viral vectors used in this study: pHR-hSyn-EGFP (Addgene #114215), pUltra-hSyn-mCherry, pHAGE-RSV-tdTomato-2A-GCaMP6f (Addgene #80317).

### High-intensity focused ultrasound

We used a custom-built focused ultrasound machine mounted on a stereotactic frame as previously described.^32^ In brief, a 6.4 cm 510 kHz transducer (H107 & Y107 Sonic Concepts) was attached within a 3D-printed coupling cone filled with degassed water. An Agilent 33220A waveform generator drove a Henry Electronics 50 watt amp and was monitored using a PVDF hydrophone (RP Acoustic, RP24l) and an oscilloscope in FFT mode (Tektronics TDS3014B). Assuming 0% attenuation, the acoustic pressure generated by this apparatus was calculated using the formula *P* = *V/M*(*f*), where *P* is the acoustic pressure, *V* is the input voltage (V, volts), and *M*(*f*) is the sensitivity of the hydrophone as a function of frequency (150 mV/MPa for the hydrophone used here).

### Cryopreservation

Organoids were fixed in 4% paraformaldehyde (VWR) for 1 hr at 4°C, washed with PBS, and dehydrated in 30% sucrose overnight. The following day, organoids were embedded in Tissue-Tek O.C.T. Compound (Sakura) and frozen in a dry ice-ethanol bath. Frozen tissue blocks were then processed into 14-μm sections using the Leica CM3050 S cryostat (Leica Biosystems) and adhered onto Superfrost Plus Microscope slides (Fisher) and stored at −80°C until use.

### Immunofluorescent staining

Tissue sections were rehydrated in 1X Tris Buffered Saline (TBS; Sigma) containing 0.1% tween-20 (TBST), permeabilized for 15 min at room temperature in TBS + 0.1% triton X-100, blocked with TBST + 5% fetal bovine serum for 30 min at room temperature, and stained overnight with primary antibodies diluted in blocking buffer. The following day, slides were washed with TBST and incubated with Alexa Fluor secondary antibodies and DAPI (Life Technologies) for 1 hr. Slides were washed 3 times with TBST and coverslips were mounted using Vectashield (Vector Laboratories). MitoTracker Red CMXRos was used according to the manufacturer’s protocol, 1h prior to fixation.

### Immunohistochemistry

Whole organoids were fixed in 10% neutral buffered formalin for 30 minutes and embedded in paraffin and sectioned at 7 μm. Antigen retrieval was performed using Tris/EDTA buffer, ph=8-8.5 for 1 hour. Immunohistochemical staining was performed using a Ventana Benchmark XT.

### Microscopy

A Zeiss AxioZoom.v16 wide-field fluorescent microscope was used for longitudinal imaging of *SYN1*::eGFP-labelled neurons. For TDP-43 imaging, a Zeiss LSM 800: AxioObserver.M2 upright confocal microscope was used with a 20X or 63X objective. To image eGFP coverage, a Leica Thunder widefield microscope coupled with computational clearing was used.

### Image analysis

#### TDP-43 & Tau

Mean fluorescence intensity of phosphorylated TDP-43 (S409/410) was measured in ImageJ and normalized to organoid area.

#### Nuclear Pores

Nuclear pores were imaged through Z-stacks spanning whole nuclei using a Zeiss LSM 800 confocal microscope. Huygens Essential (Scientific Volume Imaging (SVI)) was used for deconvolution. Imaris (BitPlane) was used to quantify nuclear pore volume and counts. Neuronal nuclei were selected by creating a 3D mask out of the surface of each neuron and excluding any signal outside of that mask. Imaris “spots” function was then used to find and measure Nup98 complexes based on the radius of the fluorescent signal. The volumes were then recorded as well as the overall count per neuron.

### Live imaging of SYN1::eGFP neurons

Cerebral organoids were transduced with lentivirus encoding *SYN1*::eGFP for 5 days and immobilized in Matrigel (VWR) prior to experimental use. Continuous Z-stacks spanning 150 μM from the organoid surface were captured daily using a Zeiss AxioZoom.v16 wide-field upright fluorescent microscope. All planes were collapsed into a single image by extended depth of focus using the Zen Pro software. Postprocessing and image alignments were performed using ImageJ software (NIH).

### RNA sequencing

Organoids were injured by HIFU and collected for RNAseq at the indicated timepoints post-injury. Supernatant was removed and organoids were lysed in RLT buffer (Qiagen). Messenger RNA was extracted using NEBNext Poly(A) mRNA Magenetic Isolation Module according to the manufacturer’s protocol, and 3’ RNA-Seq library were prepared using the 3’-Digital Gene Expression RNAseq Library Kit (Amaryllis Nucleics). Libraries were sequenced on an Illumina NextSeq 500 machine (10-25M reads, single-end, 80 bp).

RNA sequencing output was aligned to the GRCh38/hg38 reference genome using STAR alignment (STAR 2.5.3a) in PartekFlow (Partek).^64^ Genes were annotated using the GENCODE29_v2 comprehensive gene annotation. Post-alignment quality control was performed in PartekFlow. To identify differentially expressed genes, the DESeq2 (v3.5) package in PartekFlow was used to estimate dispersion and size factors, fit the data to a local model, and generate differential statistics using the Wald hypothesis test. Ingenuity Pathway Analysis (IPA, Qiagen) were used to perform downstream analysis using differential genes.

### Calcium imaging

Concentrated GCaMP6 lentivirus was transduced into organoid cultures with 1:1000 polybrene for 24 hours. GCaMP6 activity was measured by continuous imaging on a Zeiss Axiozoom.v16 for two minutes at 24 frames per second. Change in fluorescence was calculated by measuring peak fluorescent intensity per frame normalized to background fluorescence.

### Tau pThr231/total Tau ELISA measurements

Total and phosphorylated (Thr231) Tau concentrations were quantitatively measured using a 96-well electrochemiluminescence-linked immunoassay (Meso Scale Discovery, K15121D) using manufacturer protocols. Supernatant was collected from organoid culture medium and Tau/pTau concentrations were calculated after normalization to a standard curve.

### Statistical Analysis

Analysis was performed using the statistical package Prism (GraphPad Prism Version 8.4.2). Statistical analysis of neuron survival experiments was conducted using a two-sided log-rank test. For each condition, survival data from 100 *SYN1*::eGFP neurons were randomly selected and used to generate a survival curve. For all other experiments, differences between two groups were calculated using a twotailed Student’s *t*-test (when data was normally distributed) or two-sided Mann-Whitney test to compare ranks if data was not normally distributed. Differences between three or more groups were analyzed by One-way ANOVA with Tukey correction for multiple testing. Significance was assessed by *P* < 0.05. Error bars represent s.e.m. unless otherwise stated.

### Additional reagent information

The following antibodies were utilized in this manuscript: Chicken anti-MAP2 (1:10,000, Abcam; ab5392;). Rat anti-CTIP2 (1:200, Sigma; MABE1045). Mouse anti-SATB2 (1:200, Abcam; ab51502). Rabbit anti-GFAP (1:2000, Abcam; ab7260). Rabbit anti-phosphorylated TDP-43 S409/410 (1:200, Proteintech; 22309-1-AP). Mouse anti-AT8 (1:1000, Invitrogen; MN1020). Chicken anti-GFP (1:1000, Aves; GFP-1010). Rabbit anti-NUP98 (1:500, Cell Signaling;C39A3). PHF1 (1:1000) was a gift from Peter Davies.

